# Flux balance analysis of microbial communities from metabolic strategies: composition, cross-feeding and ecological service

**DOI:** 10.64898/2026.08.20.745959

**Authors:** Timothy Paez-Watson, Maria Suarez-Diez, Frank J Bruggeman

**Affiliations:** Systems Biology Lab, A-Life, AIMMS, VU University, De Boelelaan 1087, 1081 HV, Amsterdam, The Netherlands; Holomicrobiome Innovation Institute, The Netherlands; Laboratory of Systems and Synthetic Biology, Wageningen University & Research, Stippeneng 4, 6708 WE, Wageningen, The Netherlands

**Keywords:** flux balance analysis, microbial community, macrochemical equation

## Abstract

Microorganisms interact through the exchange of metabolites and competition for shared substrates, and this metabolic coupling shapes the composition and function of microbial communities. Community flux balance analysis (cFBA) can predict such behaviour – the maximum community growth rate, the metabolic fluxes and the relative abundances of the species – from stoichiometric models of their metabolism, but existing formulations are either complex and hard to scale as communities grow or cannot predict optimal growth rates. Here we present a physiology-based formulation of cFBA in which each species’ metabolism is reduced to a few macrochemical equations, one for each “metabolic mode” the species can use, and the whole community is then solved as a single linear program. From this, the method predicts the optimal composition of the community, its maximum growth rate, the metabolites exchanged between the species, and the net conversion the community carries out as a whole – its ecological service. This reduction makes it far simpler to build and solve models of larger communities. We illustrate the approach on a two-species synergistic community that can be verified by hand, apply it to a five-member anaerobic digestion community, and use it to predict the metabolic interactions of a genome-scale syngas-fermenting coculture. Characterising these communities at their optimal steady states, we show that each species is driven to a distinct metabolic strategy. We discuss the method both as a practical tool for larger microbial communities and as a means of uncovering the ecological principles that govern them.

## Introduction

Microbial communities are omnipresent. They drive the cycling of chemical elements [1], support organismal health (for example, by supplying essential vitamins in gut microbiomes [2]), enable bioremediation such as wastewater treatment, and underpin fermentative food biotechnology [3]. They also hold great promise for future biotechnology, where microbial community processes could replace conventional chemical synthesis [4, 5]. Because all of these functions rest on the net metabolic activity of the community, understanding them—so as to protect communities from pollution, antibiotics and climate change, or to exploit them in applications—calls for quantitative methods that describe community metabolism.

Such methods should answer questions such as: What metabolism does each species express when it coexists with others? How do different species compete or cooperate? What is the emergent net conversion carried out by the community as a whole, and what sets the relative abundances of its members and the striking biodiversity of some communities? Such questions are hard to answer directly: extrapolation from (meta)genomic data reveals which species are present and which metabolisms they could in principle perform [6], but not the metabolic state the community actually adopts, and even measured community fluxes do not disclose each species’ contribution, because many species consume and produce the same compounds [7].

Flux balance analysis (FBA) offers a route to these answers. Pioneered by Small & Fell [8] and Varma & Palsson [9] in the 1980s and developed continually since [10], FBA assumes that a species’ metabolism operates at steady state and predicts its flux distribution from the stoichiometry of its (genome-scale) metabolic network [11] together with an optimisation objective, usually growth-rate maximisation[12]. Because both the flux-balance constraints and the objective are linear in the fluxes, the problem is a linear program that solves quickly even for networks of thousands of reactions, and it can be applied to time-varying environments [13].

However, extension of FBA from one species to a community raises three genuine difficulties (described in detail in the Supplementary Material). First, the fluxes of FBA are, generally, specific fluxes, expressed per unit of biomass, so the net conversion a species contributes to the community is its specific flux multiplied by the amount of that species’ biomass– each flux must be weighted by a biomass amount. Second, those biomass amounts, and hence the relative abundances of the members, are themselves unknowns of the problem, and must therefore enter the optimisation as variables rather than as fixed inputs. Third, in classical FBA for a single species the evolutionary objective of metabolic optimisation is comparatively easy to define—the growth rate—whereas in a community it is not. This difficulty arises because natural selection acts at the level of individual species and not at the level of the community as a whole: the properties of a community are the outcome of selection on its member species, constrained by their interactions [14]. This has caused some confusion in the literature and has led to computational methods that impose, at least indirectly, community-level objectives, which are not in line with basic evolutionary theory.

Several methods address these difficulties in different ways that have been systematically evaluated for both software quality and predictive accuracy [15, 16] (Table 1). Community FBA (cFBA) predicts species abundances, but couples them through a bilinear program that scales poorly with community size [17]. OptCom uses multi-level optimisation [18], whereas SteadyCom imposes a common growth rate [19]. RedCom reduces complex metabolic models to elementary flux vectors but removes bilinearity by fixing the community growth rate [20].

**Table 1.**
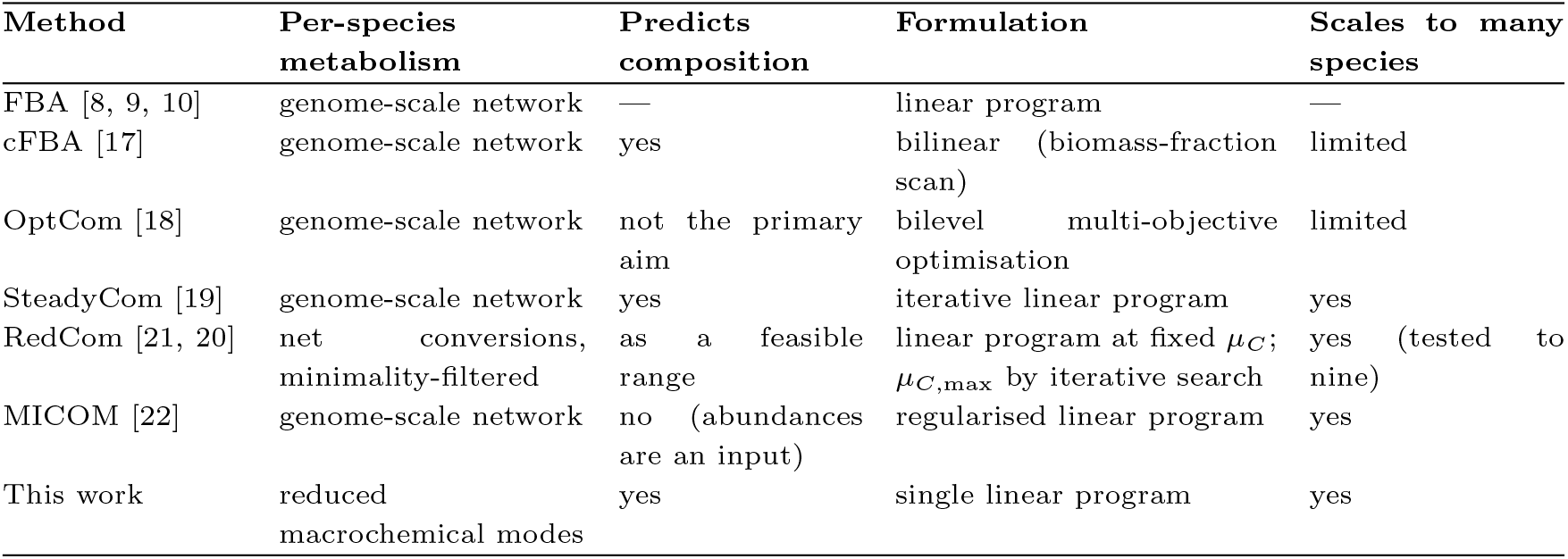
Steady-state flux balance methods for single species and microbial communities; dynamic and spatiotemporal extensions are reviewed elsewhere [15]. Most community methods operate on genome-scale metabolic networks of every member. RedCom likewise works from reduced net conversions, but retains a species-level objective and a minimality criterion on the exchange fluxes; the method presented here reduces each species to a few macrochemical modes and solves the whole community as a single linear program, while still predicting its composition.

| Method | Per-species metabolism | Predicts composition | Formulation | Scales to many species |
| --- | --- | --- | --- | --- |
| FBA [8, 9, 10] | genome-scale network | — | linear program | — |
| cFBA [17] | genome-scale network | yes | bilinear (biomass-fraction scan) | limited |
| OptCom [18] | genome-scale network | not the primary aim | bilevel multi-objective optimisation | limited |
| SteadyCom [19] | genome-scale network | yes | iterative linear program | yes |
| RedCom [21, 20] | net conversions, minimality-filtered | as a feasible range | linear program at fixed $\mu_C$; $\mu_{C,\max}$ by iterative search | yes (tested to nine) |
| MICOM [22] | genome-scale network | no (abundances are an input) | regularised linear program | yes |
| This work | reduced macrochemical modes | yes | single linear program | yes |

We recently developed a stoichiometric theory of microbial communities in which each species is described by charge- and element-balanced macrochemical reactions, and showed that steady-state cross-feeding between such reactions sets the relative abundances of the species and the net conversion carried out by the community [23]. Here we turn that description into a computational method: a physiology-based formulation of cFBA that represents each species’ metabolism as a small set of macrochemical reactions—one per metabolic mode [24]—and couples them into a single linear program. This reduction preserves what cFBA predicts (community composition, growth rate, metabolic exchanges, and the community’s emergent ecological service) while remaining transparent enough to analyse by hand and simple enough to scale. We first introduce the macrochemical formulation and its linear program on a two-species community, and then apply it to a five-member anaerobic digestion community. Finally, we apply the method to a genome-scale syngas-fermenting coculture, where it predicts the metabolic strategy that each species adopts.

## Materials and methods

The method itself—macrochemical modes, the flux balances, the weight variables and the linear program—is developed in the Results; here we only record the models, data and software used. Full derivations are given in the Supplementary Material.

### Models and data

The toy community consists of two hypothetical organisms, *P* and *Q*, with small hand-built internal networks sharing ATP and NADH; the medium supplies glucose and N_2_ at bounded rates. The anaerobic digestion community comprises five species (*C. butyricum, M. maripaludis, D. vulgaris, D. multivorans* and *M. barkeri*), each represented by a single macrochemical reaction and together involving nine external metabolites. These macrochemical equations were taken directly from our accompanying stoichiometric theory of microbial communities [23], where they were derived from the known growth physiology of each species. The syngas co-culture uses the genome-scale reconstructions of *C. autoethanogenum* (iCLAU786) and *C. kluyveri* (iCKL708) from Benito-Vaquerizo et al. [25]; a few curation fixes to the deposited models (Supplementary Material) were applied before use.

### Mode extraction

Each metabolic mode is one macrochemical reaction, which can be obtained in several complementary ways: (i) inferred from experimental data [26], (ii) chemical conservation with thermodynamic relations [27, 28];, or (iii) computed from a genome scale model using steady-state methods [10]. Toy modes come from the hand-built networks; syngas modes are extracted from the two genome-scale reconstructions. For the method to work, every mode must be tied to growth, since the weight of a mode in the community program is expressed per unit of biomass formed. Some metabolic activities a species can perform yield no growth on their own; such an activity is captured by imposing a small growth component when the mode is extracted, so that it enters the community program as a normalisable macrochemical reaction. The Supplementary Material describes this construction and the choice of the growth fraction in full.

### Community optimisation and software

The community linear programs were solved with a simplex method, so that the returned optimum is a mixture of the fewest elementary flux modes. Throughout the main text the community problem is a single linear program in which the community growth rate is maximised; no growth rate is imposed externally. Extensions of the formulation that include non-growth-associated maintenance are given in the Supplementary Material and are not used here. Genome-scale models were handled with COBRApy [29] and all linear programs were solved with a simplex solver. Code and models to reproduce the results are available at [repository URL].

## Results

### A microbial community as a single linear program

We describe each species not by its complete metabolic network but by a small number of *macrochemical reactions*: equations that summarize, per unit of biomass formed, the net nutrients a species consumes and the products it excretes when its internal metabolism runs at steady state [24, 23]. Each macrochemical reaction is one *metabolic mode* of the species, and a species that can switch strategies is given several modes among which the community optimisation is free to mix. The internal metabolites and cofactors are balanced in the reduction, so the community problem does not have intracellular variables.

We develop the method on the minimal cross-feeding community that was used to introduce cFBA in the first place [17], so that our predictions can be compared directly with the original full-stoichiometry formulation (Figure 1). The organism *P* is the only member that can use glucose and the organism *Q* the only one that can fix N_2_; neither grows alone in the medium, so the pair is an obligate mutualism. Giving *P* two modes (respire glucose, *P*_1_, or ferment part of it to succinate, *P*_2_) and *Q* two (grow self-sufficiently, *Q*_1_, or over-fix nitrogen and export ammonia, *Q*_2_) yields four macrochemical reactions, each written per unit of biomass formed (Figure 1B). Only *P*_2_ secretes succinate and only *Q*_2_ secretes ammonia, so these two cross-feeds are what couple the species.

**Fig. 1.**
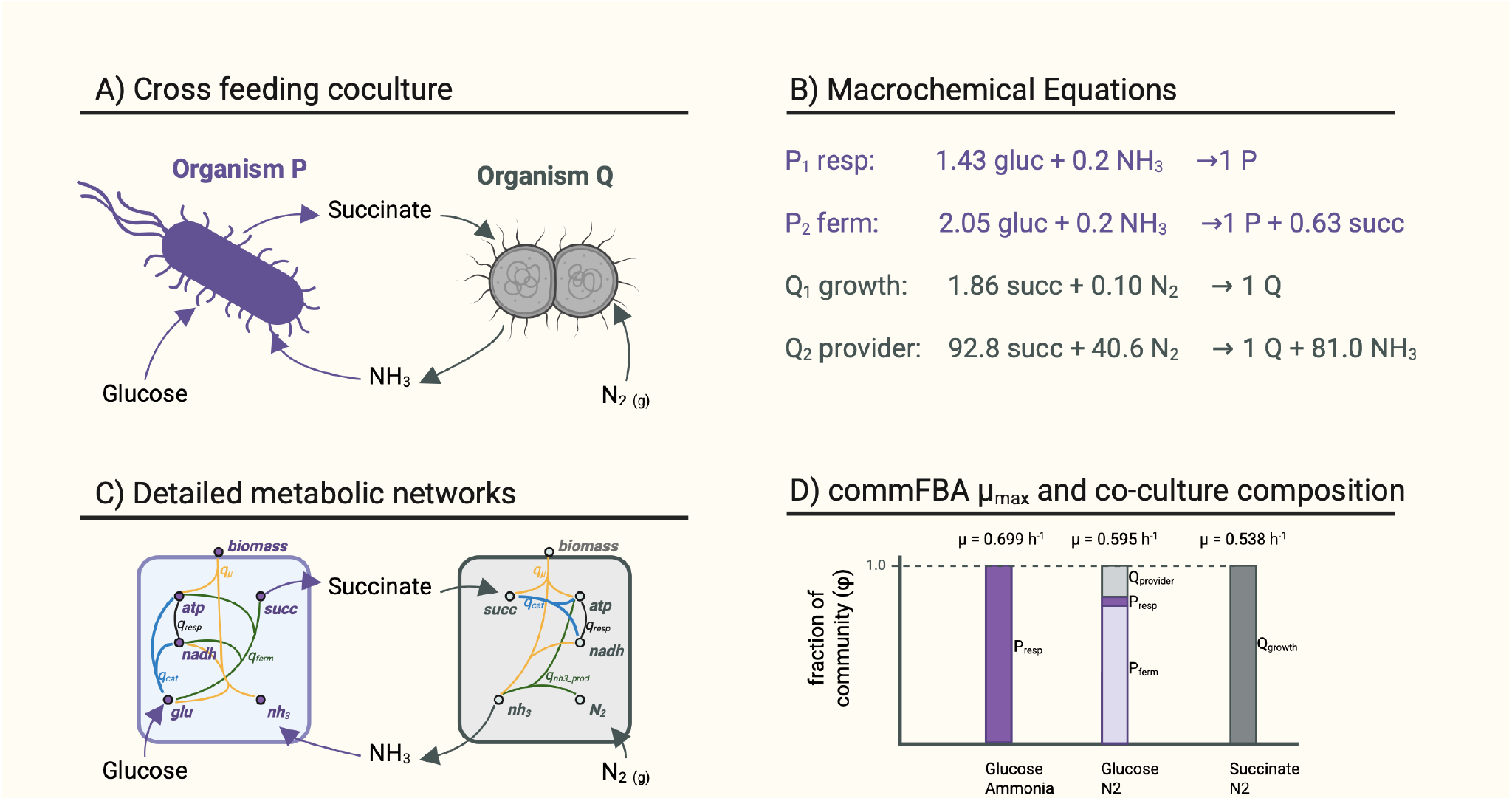
Macrochemical community FBA of a two-member cross-feeding co-culture. **(A)** The mutualism: *P* uses glucose and can secrete succinate; *Q* fixes N_2_ and can supply ammonia. Neither grows alone on the medium. **(B)** The four macrochemical reactions, one per metabolic mode, normalised per unit of biomass formed; cofactors are balanced internally and do not appear. **(C)** The small internal networks from which the equations in (B) are obtained by single-organism FBA and normalisation by growth. **(D)** Community growth rate *µ*_*C*_ and composition *ϕ* predicted by the single linear program (10) for three environments; composition, mode usage and growth rate are all outputs, only the medium bounds differ.

In this toy-model, *Q* can fix *N*_2_ and generate *NH*_3_ using the energy from succinate oxidation, without a strict links to biomass growth. However, our approach requires each *metabolic mode* to contribute to each species’s biomass growth. In such cases, a mode is extracted at a small imposed growth rate (i.e. 5 % of known *µ*_*max*_, f=0.05) which yields apparently large stoichiometric coefficients once normalized per unit biomass produced (Figure 1B). All other modes of this toy model (*P*_1_, *P*_2_ & *Q*_1_) are growth-coupled. The solution used for *Q*_2_ is required whenever a mode is disconnected from growth.

For an external metabolite *k*, steady state requires that its total production and consumption across the community balance the net exchange with the environment,

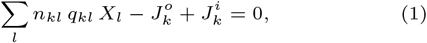

where

*q*_*kl*_ specific rate of metabolite *k* in metabolic mode *l* (mmol gDW^−1^ h^−1^);

*n*_*kl*_ *±*1, for production or consumption;

*X*_*l*_ biomass present in mode *l* (gDW);

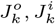 non-negative out- and inflows with the environment (mmol h^−1^).

Each specific rate must multiply a biomass amount because only then can the flow *q*_*kl*_*X*_*l*_ be balanced against the environmental exchange. This is the first difficulty named in the Introduction, and it is what makes (1) bilinear: both *q*_*kl*_ and *X*_*l*_ are unknown, and they appear multiplied together.

The bilinearity can be removed. Each mode is a fixed conversion, so its specific rates are not independent of its growth rate: doubling how fast a species grows in a given mode doubles every uptake and secretion rate in that mode. All the specific rates of mode *l* can therefore be written as

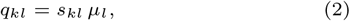

where

*s*_*kl*_ stoichiometric coefficient of metabolite *k* in that mode’s macrochemical reaction;

*µ*_*l*_ growth rate the species achieves in mode *l* (h^−1^).

The unknown product *q*_*kl*_*X*_*l*_ then becomes *s*_*kl*_ *µ*_*l*_*X*_*l*_, in which only the combination *µ*_*l*_*X*_*l*_ is unknown.

Dividing each balance (1) by the total community biomass *X*_tot_ = ∑_*l*_ *X*_*l*_, where *l* runs over all modes, collects this combination into a single variable, the *weight*

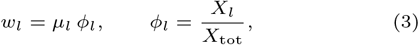

where

*ϕ*_*l*_ fraction of the community biomass in mode *l*;

*w*_*l*_ weight of mode *l* (h^−1^).

Every balance is then linear in the weights,

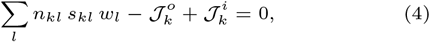

where the environmental exchanges per unit of total community biomass are

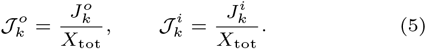

For the toy community the signed coefficients *n*_*kl*_*s*_*kl*_ are read directly from Figure 1B, giving one balance for each exchanged metabolite,

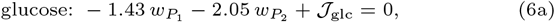

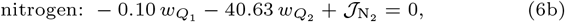

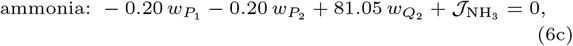

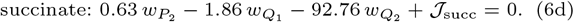

At a stable steady state the community grows with a fixed composition: the biomass fraction of every present mode, *ϕ*_*l*_ = *X*_*l*_*/X*_tot_, stays constant in time. A constant fraction has zero logarithmic derivative, and since 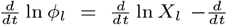 ln *X*_tot_, this forces every present mode to grow at the same rate as the community as a whole,

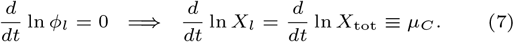

We call *µ*_*C*_ the specific (per-capita) community growth rate. This common rate is forced, not assumed: a mode growing more slowly than *µ*_*C*_ would lose biomass share every generation until it vanished, so only compositions in which every present mode shares the single rate *µ*_*C*_ can persist. Maximising *µ*_*C*_ is thus not a community-level objective imposed by hand, it is what selection on the individual species produces, each growing as fast as its environment and its partners allow.

Because every present mode grows at *µ*_*C*_, its weight is *w*_*l*_ = *µ*_*C*_ *ϕ*_*l*_, and since the biomass fractions sum to one,

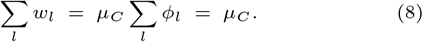

The sum of the weights *is* the common growth rate, so maximising ∑ _*l*_ *w*_*l*_ is the same as maximising *µ*_*C*_ ; the community problem inherits the objective of single-species FBA rather than needing a new one. Note that (8) holds for any value of *µ*_*C*_, so the growth rate need not be fixed in advance and searched over, as it must be in formulations that linearise the problem by holding it constant [19, 20]. The community is then solved by the single linear program

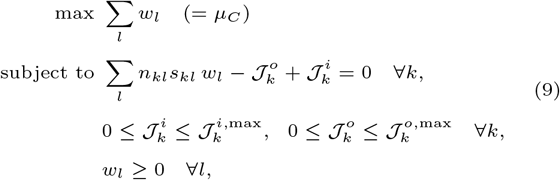

where the upper bounds 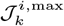 and 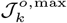 on the exchanges encode the medium and the reaction directions. For the toy community in the synergistic environment (glucose and N_2_ supplied, ammonia and succinate exchanged) the program reads

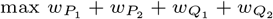

subject to the four balances (6a)–(6d),

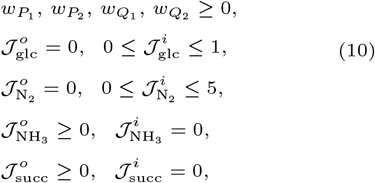

with all exchange bounds in mmol gDW^−1^ h^−1^ (per unit of total community biomass).

The internal cross-feeds are what make the synergy obligate, and this can be read directly off the balances. Ammonia has no external source 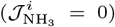, so in (6c) the consumption by *P* (the 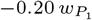 and −0.20 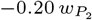 terms) can only be met by production from *Q*_2_ (the +81.05 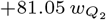 term); hence 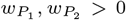 requires 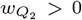. Symmetrically, succinate has no external source 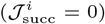, so in (6d) the consumption by *Q* (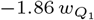 and 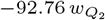) can only be met by production from 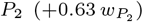; hence 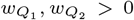 requires 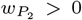. The mutualism is therefore imposed by the constraints rather than assumed, and maximising *µ*_*C*_ sets the balance between fermentation by *P* and nitrogen fixation by *Q*.

Relative abundances follow from the optimal weights as *ϕ*_*l*_ = *w*_*l*_*/µ*_*C*_ ; they are an *output* of the program, not an input. Solving (10) reproduces the result obtained for this same community by the original, full-stoichiometry cFBA of Khandelwal et al. [17] to within 5 × 10^−4^ (*µ*_*C*_ = 0.605 versus 0.6047 h^−1^), but as one linear program rather than a scan over biomass ratios. Changing only the medium bounds returns qualitatively different communities (Figure 1D): supplying ammonia directly makes *Q* redundant and lets *P* respire (*µ*_*C*_ = 0.70 h^−1^); glucose with N_2_ alone restores the mutualism at a *P* :*Q* ratio near 5:1 (*µ*_*C*_ = 0.60 h^−1^); and succinate in place of glucose removes *P*, so *Q* grows self-sufficiently (*µ*_*C*_ = 0.54 h^−1^). Composition, mode usage and growth rate are all outputs of the same program.

### Scaling to a multispecies community: anaerobic digestion

We applied our method to anaerobic digestion, in which a community cross-feeds short-chain fatty acids and alcohols to turn organic matter into methane and carbon dioxide [30, 31, 32], and which has itself been a recurring test case for reduced community models [20]. Here glucose is digested to methane and carbon dioxide by a five-member community—*C. butyricum* (cb), *M. maripaludis* (mm), *D. vulgaris* (dv), *D. multivorans* (dm) and *M. barkeri* (mb)—that imports glucose, ammonium and sulfate, exports methane, carbon dioxide and bisulphide, and cross-feeds acetate, hydrogen and butyrate internally (Figure 2A). Each species is a single macrochemical reaction, giving nine external-metabolite balances and one linear program, exactly as for the toy community.

**Fig. 2.**
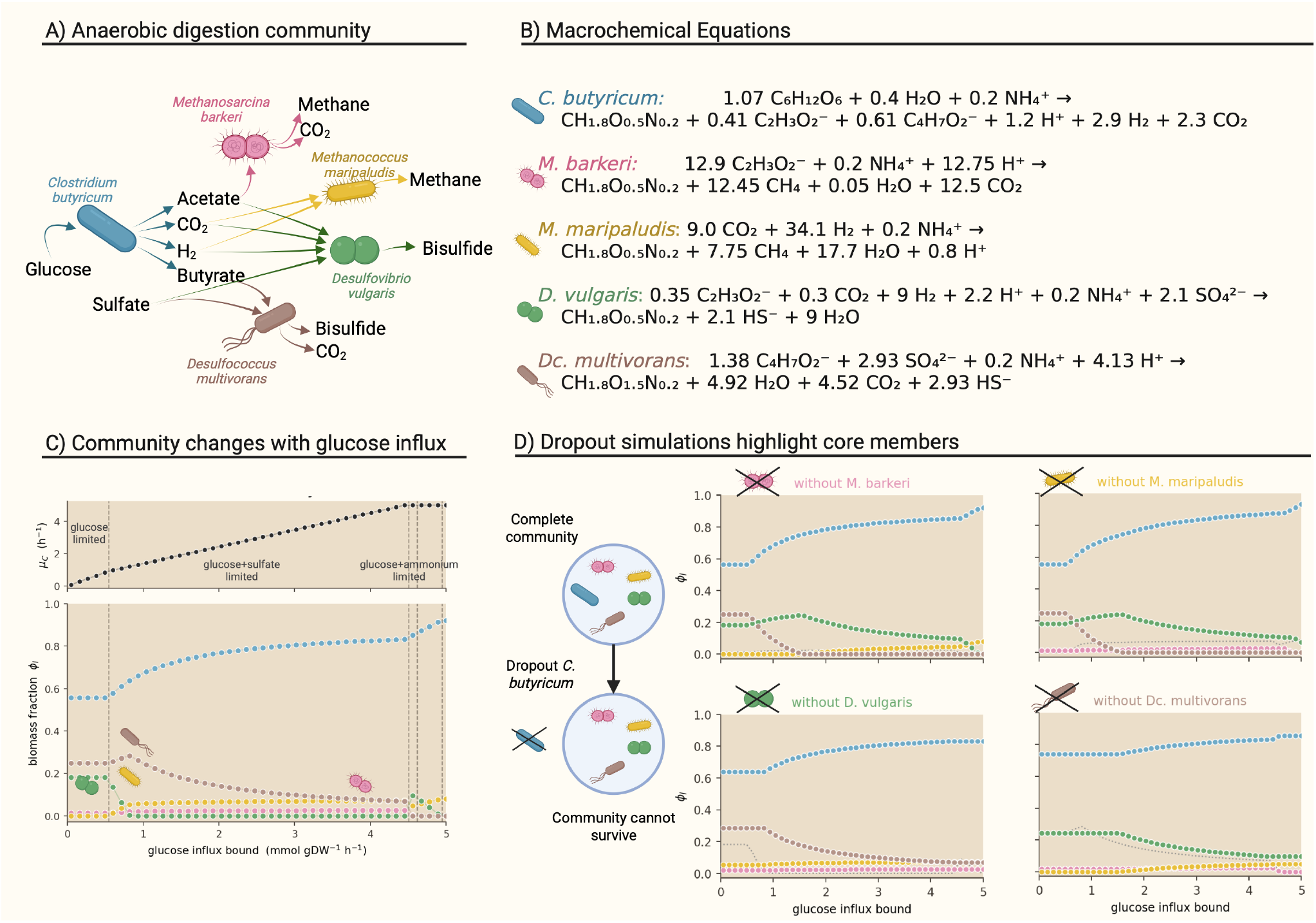
A five-member anaerobic digestion community. **(A)** The community imports glucose, ammonium and sulfate and cross-feeds acetate, hydrogen and butyrate en route to methane and carbon dioxide; each species is one macrochemical reaction. **(B)** Community growth rate and composition *ϕ* as a function of the glucose influx; composition and the net conversion change between flux-limitation regimes. **(C)** Single-species knockouts distinguish a keystone (*C. butyricum*) from a substitutable member (*M. maripaludis*).

Scanning the glucose influx traces how both the community growth rate and its composition respond to resource supply (Figure 2B). The optimum passes through distinct regimes set by which environmental supply is limiting, and the microbiome composition changes with it: all five species coexist only in a narrow window, while at higher glucose *D. vulgaris* is excluded as sulfate becomes limiting.

At each steady state the community catalyses a net conversion, which we call its *ecological service*. Once the optimal weights are known, every internal exchange cancels by construction: whatever one species secretes and another consumes appears in (4) with equal and opposite sign. What survives is the net traffic with the environment, ∑_*l*_ *n*_*kl*_*s*_*kl*_*w*_*l*_ for each metabolite *k*. A community of five species is thereby summarised by one equation, and that equation is a prediction. At a glucose uptake bound of one, where glucose and sulfate both bind and four of the five species are present, the ecological service is

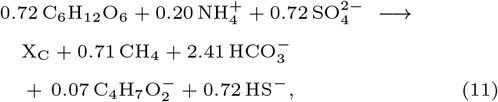

with X_C_ one unit of community biomass. The service is not a fixed property of the community but of the community in an environment. Raising only the glucose supply, and leaving the ammonium and sulfate supplies exactly as they were, gives a visibly different equation,

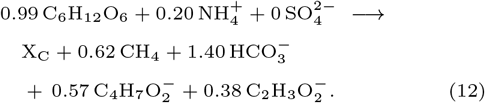

With ample glucose, ammonium is the binding constraint, and since every species requires the same 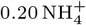 per unit of biomass the nitrogen supply alone fixes *µ*_*C*_ . Sulfate reduction, in this case, yields no additional growth, hence both sulfate reducers are absent from the optimum.

Single-species knockouts separate two kinds of members. Removing *C. butyricum*, the sole point of glucose entry and hence part of every mode, collapses the community; removing the hydrogen-consuming specialist *M. maripaludis* costs almost no growth, its role being taken over by *D. vulgaris* (Figure 2C). Neither outcome is predictable from abundance alone.

### Using genome-scale models: a syngas-fermenting co-culture

The modes need not come from hand-built physiology; they can be read off genome-scale reconstructions. We reduced the two-member syngas co-culture of the acetogen, *Clostridium autoethanogenum* (iCLAU786) and the chain elongator *Clostridium kluyveri* (iCKL708) studied by Benito-Vaquerizo et al. [25] —together 2064 reactions and 1823 metabolites—to eight macrochemical modes, five for the acetogen and three for the chain elongator (Figure 3A). The extraction follows the same logic as for the toy community, applied one organism at a time. Each substrate combination the organism can grow on, and each qualitatively distinct product it can make from them, is imposed in turn on its genome-scale model; the resulting flux distribution is normalised by growth and read off as one balanced macrochemical equation. The set obtained this way spans the main strategies each organism has available, and no composition, exchange or mode usage is fixed: all of them are returned by the community program.

**Fig. 3.**
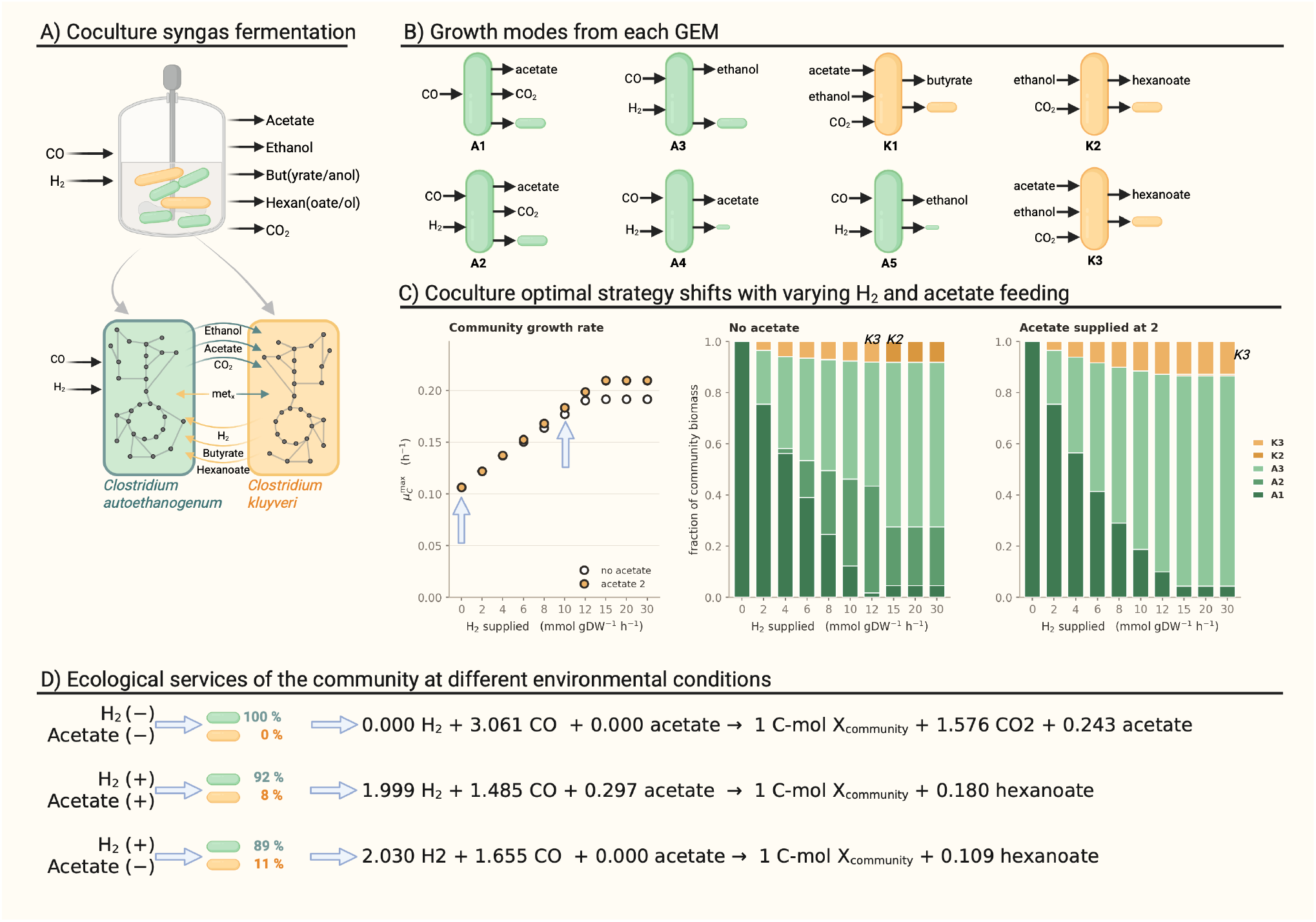
A genome-scale syngas co-culture reduced to macrochemical modes and solved for maximum community growth. **(A)** The co-culture: *Clostridium autoethanogenum* (an acetogen, green) and *Clostridium kluyveri* (a chain elongator, orange) grown together on syngas in a chemostat fed with CO and H_2_, releasing acetate, ethanol, butyrate/butanol, hexanoate/hexanol and CO_2_. The acetogen fixes CO and H_2_ and hands ethanol, acetate and CO_2_ to the chain elongator, which condenses these into longer-chain acids and can return H_2_; the pair was characterised experimentally and modelled by Benito-Vaquerizo et al. [25]. **(B)** The two genome-scale reconstructions (iCLAU786 and iCKL708; together 2064 reactions and 1823 metabolites) are reduced to eight macrochemical modes—five for the acetogen (*A*_1_–*A*_5_) and three for the chain elongator (*K*_1_–*K*_3_)—each a substrate-to-product conversion normalised per unit of biomass formed. Modes *A*_4_ and *A*_5_ represent overflow to acetate and ethanol at low growth, and are drawn with a small biomass product to reflect this; the stoichiometric coefficients of all modes are listed in the Supplementary Material and omitted here for clarity. **(C)** Optimal strategy as the H_2_ supply is scanned, without an acetate feed and with acetate supplied at 2 mmol gDW^−1^ h^−1^. Left: maximum community growth rate 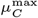 for the two feeds (open: no acetate; filled: acetate at 2). Middle and right: the corresponding community composition, given as the biomass fraction in each mode. Without H_2_ the optimum is a monoculture of the acetogen; as H_2_ rises the acetogen switches from acetate production (*A*_1_/*A*_2_) to ethanol production (*A*_3_) and the chain elongator (*K*_2_/*K*_3_) is recruited, so the two species coexist and 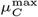 increases until carbon becomes limiting. **(D)** The ecological service—the net macrochemical conversion carried out per C-mol of community biomass—for three feed conditions from (C), each labelled with the optimal acetogen:chain-elongator composition (green:orange). Without H_2_ the acetogen grows alone and the community excretes CO_2_ and acetate; supplying H_2_, with or without an acetate feed, brings in the chain elongator and hexanoate becomes the only organic product leaving the system.

Maximising the community growth rate across a hydrogen supply highlights the interdependence and conditions for this coculture. Without hydrogen the optimum is a monoculture: the acetogen converts carbon monoxide to biomass more efficiently on its own than the community can by routing carbon through a partner. Supplying hydrogen changes the balance. The extra electrons let the acetogen switch from making acetate to making ethanol, and ethanol is what the chain elongator needs to run reverse *β*-oxidation and build longer carbon chains [33]; coexistence becomes the optimum, with the partner’s share and the community growth rate both rising until carbon monoxide becomes limiting (Figure 3B,C). The two-carbon intermediates are then produced and consumed entirely inside the community and hexanoate is the only organic product that leaves this system.

Feeding acetate sharpens the same mechanism. It does nothing without hydrogen, since the chain elongator cannot use acetate without ethanol, but once hydrogen is present it raises both the growth rate and the share of the chain elongator by relieving the acetogen of having to make the two-carbon currency itself, so that it commits further to ethanol. Genome-scale detail is thus not needed to say which organisms coexist, why, and which conversion each of them runs.

## Discussion

Reducing each species to a handful of macrochemical modes and rewriting the community balances in terms of the mode weights turns community flux balance analysis into a single linear program whose solution includes the community composition. The same construction handled a hand-built two-member syntrophy, a five-member anaerobic digestion community, and a genome-scale syngas co-culture, in every case returning relative abundances, metabolic exchanges and the community’s net conversion as outputs. Because the reduction removes all intracellular variables, the problem stays small and transparent: the syngas co-culture collapses from 2064 reactions to eight equations while still predicting when the two organisms coexist and which conversion each of them runs.

It is worth being precise about what the reduction itself buys, since reduced community models are not new [21, 20]. The gain is not smaller models but a linear one. Because (8) makes the sum of the weights identically equal to the shared growth rate, the balanced-growth condition is satisfied by construction and *µ*_*C*_ is the objective value of a single linear program, rather than a parameter that must be fixed and then searched over. That also settles the objective. Reduced formulations have set growth-rate maximisation aside because it admits solutions in which a species overproduces for its partners at its own expense, and have replaced it with criteria of substrate efficiency [20]; such criteria are effective, but they are imposed on the community rather than derived from selection on its members. In the weight variables no such criterion is needed, because maximising ∑_*l*_ *w*_*l*_ *is* each species growing as fast as its environment and its partners allow. A practical consequence is numerical: bilinear community models are known to return different optima from different starting points [20], whereas a linear program has none of that behaviour.

Because composition is returned rather than supplied, the method distinguishes a function a community lacks from one it possesses but does not use at the optimum. The digestion community stops consuming sulfate entirely, and both sulfate reducers disappear, at glucose supplies where nitrogen rather than sulfate is limiting—although sulfate is still present and two members are equipped to perform it. It is simply that no allocation of biomass to those species increases the community growth rate once nitrogen sets the ceiling. A method given the relative abundances as input cannot draw this distinction, because the absences it would have to explain were never predicted in the first place.

We think the ecological service is the most useful quantity the method returns, and the one that generalises furthest beyond the systems treated here. It is a single balanced macrochemical equation describing what a community does to its surroundings: what it draws in, what it releases, and in what proportions. For microbiome research this is the quantity that is usually wanted and rarely obtained: sequencing surveys report who is present, and metabolic reconstruction reports what each member could in principle do, but neither yields the net conversion the assembly actually carries out. Expressed per unit of community biomass, the service is directly comparable across habitats—a gut community, a bioreactor and a sediment can be placed on the same axis and asked whether they perform the same chemistry, without requiring them to share species, a host, or an experimental setting. The same construction is important for biotechnological applications making use of microbial communities, since the service represents how a microbiome transforms substrates into desired products. An engineer can predict environmental conditions that favour higher production yields in a given community. Methods simple enough to scale are what make this move from cataloguing membership to predicting and steering function practical

The service is not static, however. As the two digestion conversions show, it shifts as the strategies change in response to the environmental conditions. Truly steady state conditions are seldomly the case in natural environments where microbiomes live and act, and thus extending the prediction of ecological functions to dynamic conditiosn is an important next step to take. The natural next step is therefore to carry this reduction into a dynamic setting, using the same macrochemical simplification to keep the problem tractable while allowing the community and its service to change in time. That is an open question, and one we are currently pursuing.

## Supplementary Materials

Supplementary materials are provided with the publication of this paper. Code, models and the scripts that generate every figure and table in the main text and in this document are available at https://github.com/TP-Watson/Community-FBA-from-metabolic-strategies/tree/main.

## References

1. Paul G. Falkowski, Tom Fenchel, and Edward F. Delong. The microbial engines that drive Earth’s biogeochemical cycles. Science, 320(5879):1034–1039, 2008.

2. Jean Guy LeBlanc, Christian Milani, Graciela Savoy de Giori, Fernando Sesma, Douwe van Sinderen, and Marco Ventura. Bacteria as vitamin suppliers to their host: a gut microbiota perspective. Current Opinion in Biotechnology, 24(2):160–168, 2013.

3. Benjamin E. Wolfe and Rachel J. Dutton. Fermented foods as experimentally tractable microbial ecosystems. Cell, 161(1):49–55, 2015.

4. Nicholas S. McCarty and Rodrigo Ledesma-Amaro. Synthetic biology tools to engineer microbial communities for biotechnology. Trends in Biotechnology, 37(2):181–197, 2019.

5. Prashant Kumar, Dheeraj Chitara, Sourodip Sengupta, Paromita Banerjee, and Sachchida Nand Rai. Microbial consortia in biotechnology: applications and challenges in industrial processes. 3 Biotech, 15(11):386, 2025.

6. Eric A. Franzosa, Tiffany Hsu, Alexandra Sirota-Madi, Afrah Shafquat, Galeb Abu-Ali, Xochitl C. Morgan, and Curtis Huttenhower. Sequencing and beyond: integrating molecular ‘omics’ for microbial community profiling. Nature Reviews Microbiology, 13(6):360–372, 2015.

7. Marc G. Dumont and J. Colin Murrell. Stable isotope probing—linking microbial identity to function. Nature Reviews Microbiology, 3(6):499–504, 2005.

8. David A. Fell and J. Rankin Small. Fat synthesis in adipose tissue: an examination of stoichiometric constraints. Biochemical Journal, 238(3):781–786, 1986.

9. Amit Varma and Bernhard O. Palsson. Stoichiometric flux balance models quantitatively predict growth and metabolic by-product secretion in wild-type Escherichia coli W3110. Applied and Environmental Microbiology, 60(10):3724–3731, 1994.

10. Jeffrey D. Orth, Ines Thiele, and Bernhard Ø. Palsson. What is flux balance analysis? Nature Biotechnology, 28(3):245–248, 2010.

11. Ines Thiele and Bernhard Ø. Palsson. A protocol for generating a high-quality genome-scale metabolic reconstruction. Nature Protocols, 5(1):93–121, 2010.

12. J.S. Edwards and B.O. Palsson. Robustness analysis of the escherichia coli metabolic network. Biotechnology Progress, 16(6):927–939, December 2000.

13. Radhakrishnan Mahadevan, Jeremy S. Edwards, and Francis J. Doyle. Dynamic flux balance analysis of diauxic growth in Escherichia coli. Biophysical Journal, 83(3):1331–1340, 2002.

14. Ali R. Zomorrodi and Daniel Segrè. Genome-driven evolutionary game theory helps understand the rise of metabolic interdependencies in microbial communities. Nature Communications, 8:1563, 2017.

15. William T. Scott, Jr., Sara Benito-Vaquerizo, Johannes Zimmermann, Djordje Bajić, Almut Heinken, Maria Suarez-Diez, and Peter J. Schaap. A structured evaluation of genome-scale constraint-based modeling tools for microbial consortia. PLoS Computational Biology, 19(8):e1011363, 2023.

16. Clémence Joseph, Haris Zafeiropoulos, Kristel Bernaerts, and Karoline Faust. Predicting microbial interactions with approaches based on flux balance analysis: an evaluation. BMC Bioinformatics, 25:36, 2024.

17. Ruchir A. Khandelwal, Brett G. Olivier, Wilfred F. M. Röling, Bas Teusink, and Frank J. Bruggeman. Community flux balance analysis for microbial consortia at balanced growth. PLoS ONE, 8(5):e64567, 2013.

18. Ali R. Zomorrodi and Costas D. Maranas. OptCom: a multi-level optimization framework for the metabolic modeling and analysis of microbial communities. PLoS Computational Biology, 8(2):e1002363, 2012.

19. Siu Hung Joshua Chan, Margaret N. Simons, and Costas D. Maranas. SteadyCom: predicting microbial abundances while ensuring community stability. PLoS Computational Biology, 13(5):e1005539, 2017.

20. Sabine Koch, Fabian Kohrs, Patrick Lahmann, Thomas Bissinger, Stefan Wendschuh, Dirk Benndorf, Udo Reichl, and Steffen Klamt. RedCom: a strategy for reduced metabolic modeling of complex microbial communities and its application for analyzing experimental datasets from anaerobic digestion. PLoS Computational Biology, 15(2):e1006759. 2019.

21. Reed Taffs, John E. Aston, Kristen Brileya, Zackary Jay, Christian G. Klatt, Shawn McGlynn, Natasha Mallette, Scott Montross, Robin Gerlach, William P. Inskeep, David M. Ward, and Ross P. Carlson. In silico approaches to study mass and energy flows in microbial consortia: a syntrophic case study. BMC Systems Biology, 3:114, 2009.

22. Christian Diener, Sean M. Gibbons, and Osbaldo Resendis-Antonio. MICOM: metagenome-scale modeling to infer metabolic interactions in the gut microbiota. mSystems, 5(1):e00606.–19, 2020.

23. Frank J. Bruggeman, Timothy Paez-Watson, Bas Teusink, and Robbert Kleerebezem. Stoichiometric analysis of microbial communities links function, structure, and biomass carrying capacity. The ISME Journal, page wrag133, 2026.

24. J. A. Roels. Application of macroscopic principles to microbial metabolism. Biotechnology and Bioengineering, 22(12):2457–2514, 1980.

25. Sara Benito-Vaquerizo, Martijn Diender, Ivette Parera Olm, Vitor A. P. Martins dos Santos, Peter J. Schaap, Diana Z. Sousa, and Maria Suarez-Diez. Modeling a co-culture of Clostridium autoethanogenum and Clostridium kluyveri to increase syngas conversion to medium-chain fatty-acids. Computational and Structural Biotechnology Journal, 18:3255–3266, 2020.

26. J. J. Heijnen and J. P. Van Dijken. In search of a thermodynamic description of biomass yields for the chemotrophic growth of microorganisms. Biotechnology and Bioengineering, 39(8):833–858, 1992.

27. Robbert Kleerebezem and Mark C. M. van Loosdrecht. A generalized method for thermodynamic state analysis of environmental systems. Critical Reviews in Environmental Science and Technology, 40(1):1–54, 2010.

28. Christina M. Smeaton and Philippe Van Cappellen. Gibbs energy dynamic yield method (GEDYM): Predicting microbial growth yields under energy-limiting conditions. Geochimica et Cosmochimica Acta, 241:1–16, 2018.

29. Ali Ebrahim, Joshua A. Lerman, Bernhard O. Palsson, and Daniel R. Hyduke. COBRApy: COnstraints-Based Reconstruction and Analysis for Python. BMC Systems Biology, 7:74, 2013.

30. Glen D’Souza, Shraddha Shitut, Daniel Preussger, Ghada Yousif, Silvio Waschina, and Christian Kost. Ecology and evolution of metabolic cross-feeding interactions in bacteria. Natural Product Reports, 35(5):455–488, 2018.

31. Takashi Narihiro, Masaru K. Nobu, Na-Kyung Kim, Yoichi Kamagata, and Wen-Tso Liu. The nexus of syntrophy-associated microbiota in anaerobic digestion revealed by long-term enrichment and community survey. Environmental Microbiology, 17(5):1707–1720, 2015.

32. Ling Leng, Peixian Yang, Shubham Singh, Huichuan Zhuang, Linji Xu, Wen-Hsing Chen, Jan Dolfing, Dong Li, Yan Zhang, Huiping Zeng, Wei Chu, and Po-Heng Lee. A review on the bioenergetics of anaerobic microbial metabolism close to the thermodynamic limits and its implications for digestion applications. Bioresource Technology, 247:1095–1106, 2018.

33. Paul J. Weimer and Richard A. Kohn. Impacts of ruminal microorganisms on the production of fuels: how can we intercede from the outside? Applied Microbiology and Biotechnology, 100(8):3389–3398, 2016.

