## Supplementary Materials for "Flux balance analysis of microbial communities from metabolic strategies: composition, cross-feeding and ecological service"

**S4 Removing the bilinearity** 4

**S5 The community objective** 4

**S6 Metabolic modes and the growth fraction** 5

**S8 Extension: non-growth-associated maintenance** 9

**S9 Software, data and reproducibility** 9

### S1 Scope and notation

This document provides the derivations, model definitions and control analyses supporting the main text. Sections S2–S5 develop the formulation from first principles; Section S6 defines mode extraction and reports the sensitivity of the community optimum to the growth fraction; Section S7 gives the complete stoichiometry of all three models; Section S8 describes an extension to non-growth-associated maintenance; and Section S9 records software and data availability.

The notation of the main text is used throughout. Symbols are collected in Table S1.

Table S1: Notation and units. Subscript $l$ indexes metabolic modes, $k$ external metabolites, and $i$ species. gDW denotes grams of biomass dry weight.

| Symbol | Meaning | Units | Status |
|---|---|---|---|
| $v_j$ | reaction rate in a single-species network | $\mathrm{mmol\,gDW^{-1}h^{-1}}$ | internal |
| $q_{kl}$ | specific exchange rate of metabolite $k$ in mode $l$ | $\mathrm{mmol\,gDW^{-1}h^{-1}}$ | variable |
| $s_{kl}$ | stoichiometric coefficient of $k$ in mode $l$ | $\mathrm{mmol\,gDW^{-1}}$ | constant |
| $n_{kl}$ | sign convention, $+1$ produced, $-1$ consumed | — | constant |
| $X_l$ | biomass present in mode $l$ | gDW | variable |
| $X_T$ | total community biomass, $\sum_l X_l$ | gDW | variable |
| $\phi_l$ | biomass fraction of mode $l$, $X_l/X_T$ | — | **output** |
| $\mu_l$ | growth rate achieved in mode $l$ | $\mathrm{h^{-1}}$ | variable |
| $\mu_C$ | community growth rate | $\mathrm{h^{-1}}$ | **output** |
| $w_l$ | weight of mode $l$, $\mu_l \phi_l$ | $\mathrm{h^{-1}}$ | LP variable |
| $J_k^o, J_k^i$ | extensive out- and inflow of $k$ | $\mathrm{mmol\,h^{-1}}$ | variable |
| $\mathcal{J}_k^o, \mathcal{J}_k^i$ | the same, per unit community biomass | $\mathrm{mmol\,gDW^{-1}h^{-1}}$ | LP variable |
| $f$ | growth fraction used to extract a near-growthless mode | — | parameter |

### S2 Flux balance analysis for a single species

A metabolic network of one species contains $N_R$ reactions connecting $N_M$ internal metabolites. With the stoichiometric coefficients collected in $\mathbf{N} \in \mathbb{R}^{N_M \times N_R}$ and the reaction rates in $\mathbf{v}$, the steady-state assumption on internal metabolite concentrations is

$$\mathbf{N}\mathbf{v} = \mathbf{0}. \quad \text{(S1)}$$

#### S2.1 Units

The rates in $\mathbf{v}$ are typically specific fluxes, normalised to biomass, with units $\mathrm{mmol\,gDW^{-1}h^{-1}}$. This normalisation (not always the case, but in practice very often used) is imposed by the biomass reaction, a pseudo-reaction whose stoichiometry specifies the millimoles of each precursor required to synthesise one gram of biomass. Its rate therefore has units $\mathrm{gDW\,gDW^{-1}h^{-1}} = \mathrm{h^{-1}}$ and is the specific growth rate $\mu$. Because every flux is expressed per gram of biomass, a single-species flux vector describes the activity of one gram of that species and is independent of how much biomass is present.

#### S2.2 Underdetermination and the objective

Typically $N_R > N_M$, so (S1) does not determine $\mathbf{v}$. Flux balance analysis closes the system by bounding the fluxes, $\mathbf{v}^{\min} \leq \mathbf{v} \leq \mathbf{v}^{\max}$, to encode reversibility and capacity, and by selecting the feasible flux vector that maximises a linear objective $\mathbf{c}^\top \mathbf{v}$, conventionally the growth rate. Both the constraints and the objective are linear, so the problem is a linear program.

For a single species the amount of biomass never enters the formulation. The three complications described next all follow from the fact that this ceases to be true for a community.

### S3 Three complications in a community

A community comprises $N_S$ species sharing an environment and exchanging metabolites. The first complication concerns units, the second introduces unknowns, and the third concerns the objective.

#### S3.1 Specific fluxes must be multiplied by biomass amounts

When modelling a microbial community, metabolites exist inside the cells (as in conventional FBA), but also outside the cells which are shared amongst species. Their balance depends not on the per-gram activity of a species but on its total contribution, which scales with the amount of that species present. For external metabolite $k$,

$$\sum_l n_{kl}\, q_{kl}\, X_l \;-\; J_k^o \;+\; J_k^i \;=\; 0. \tag{S2}$$

The unit algebra requires the biomass multiplier:

$$\underbrace{q_{kl}}_{\text{mmol gDW}^{-1}\text{h}^{-1}} \times \underbrace{X_l}_{\text{gDW}} = \underbrace{q_{kl}X_l}_{\text{mmol h}^{-1}}\;. \tag{S3}$$

Only the extensive flow $q_{kl}X_l$ is dimensionally commensurate with the environmental exchanges $J_k$, which are themselves extensive. In a single-species model the multiplier is absent because there is one biomass pool and the entire problem can be posed in specific units. In a community the biomass amounts convert each species' per-gram description into the common currency in which the exchanges must balance.

#### S3.2 Community composition is unknown

The multipliers $X_l$ in (S2) are not measurable parameters of the formulation but quantities to be predicted. Defining the total biomass $X_T = \sum_l X_l$ and the biomass fractions

$$\phi_l = \frac{X_l}{X_T}, \qquad \sum_l \phi_l = 1, \tag{S4}$$

the composition $(\phi_1, \ldots, \phi_{N_S})$ enters the optimisation as a vector of variables. A single-species model has no analogue.

Composition is further constrained by the meaning of steady state for a community. If the biomass fractions are constant, every species present grows at the same specific rate, and that shared rate is the community growth rate:

$$\frac{\mathrm{d}\ln X_l}{\mathrm{d}t} = \frac{\mathrm{d}\ln X_T}{\mathrm{d}t} = \mu_C \qquad \text{for all present } l. \tag{S5}$$

#### S3.3 The objective must reflect selection on species

For a single species the objective is uncontroversial: selection favours faster growth. For a community, maximising a community-level quantity is not equivalent, because natural selection acts on individual species and not on the community as a unit. Community properties are consequences of selection on the members, constrained by their interactions. Section S5 shows that under the balanced-growth condition (S5) the objective of the community problem is determined by per-species selection and need not be postulated separately.

### S4 Removing the bilinearity

The balance (S2) contains the product $q_{kl}X_l$ in which both factors are unknown, so the constraint is bilinear and the feasible set is non-convex. This is why the original cFBA formulation scans over biomass ratios rather than solving a single program.

Each mode is a fixed conversion, so its specific rates are proportional to its growth rate. Writing $q_{kl} = s_{kl}\mu_l$ with $s_{kl}$ the constant coefficient of metabolite $k$ in the macrochemical equation of mode $l$, the offending term becomes

$$q_{kl}X_l = s_{kl}\,\mu_l X_l, \tag{S6}$$

in which only the product $\mu_l X_l$ is unknown. Dividing (S2) by $X_T$ and defining the weight

$$w_l = \mu_l\,\phi_l, \tag{S7}$$

every balance becomes linear in the weights and in the normalised exchanges $\mathcal{J}_k^o = J_k^o/X_T$, $\mathcal{J}_k^i = J_k^i/X_T$:

$$\sum_l n_{kl}\,s_{kl}\,w_l - \mathcal{J}_k^o + \mathcal{J}_k^i = 0. \tag{S8}$$

The substitution is exact. No growth rate is fixed and no composition is assumed; the bilinear term is absorbed into a single variable per mode.

### S5 The community objective

We show that maximising $\sum_l w_l$ is equivalent to per-species selection, and that its optimal value is the community growth rate.

At a steady composition the biomass fractions are constant, $\mathrm{d}\phi_l/\mathrm{d}t = 0$, which by (S4) requires every present mode to grow at the rate of the total, as in (S5). The community growth rate is the abundance-weighted mean of the species' growth rates:

$$\mu_C = \frac{\mathrm{d}\ln X_T}{\mathrm{d}t} = \frac{1}{X_T}\frac{\mathrm{d}}{\mathrm{d}t}\sum_l X_l = \sum_l \phi_l \frac{\mathrm{d}\ln X_l}{\mathrm{d}t}. \tag{S9}$$

With $\mathrm{d}\ln X_l/\mathrm{d}t = \mu_l$ and the weight (S7),

$$\mu_C = \sum_l \phi_l \mu_l = \sum_l w_l. \tag{S10}$$

Three consequences follow.

**(i) The objective is not a community-level criterion.** The quantity $\sum_l w_l$ *is* the growth rate shared by every member. Maximising it is the statement that each species grows as fast as its environment and its partners permit. No efficiency criterion, altruism penalty or multi-level objective is introduced.

**(ii) Balanced growth holds by construction.** Because $\phi_l = w_l/\sum_m w_m$ is a definition, any mode with $w_l > 0$ satisfies $\mu_l = w_l/\phi_l = \sum_m w_m = \mu_C$. The coexistence condition (S5) is therefore an identity of the formulation rather than a constraint that must be imposed.

**(iii) The growth rate need not be fixed or searched for.** Relation (S10) holds for every value of $\mu_C$, so $\mu_C$ is the objective *value* of the linear program and is obtained directly. Formulations that linearise the community problem by holding $\mu_C$ constant must instead locate its maximum by an outer search over that scalar.

The complete program is

$$\begin{aligned} \max\ & \textstyle\sum_l w_l \quad (= \mu_C) \\ \text{s.t.}\ & \textstyle\sum_l n_{kl} s_{kl} w_l - \mathcal{J}_k^o + \mathcal{J}_k^i = 0 \quad \forall k, \\ & w_l \geq 0, \quad \mathcal{J}_k^o, \mathcal{J}_k^i \geq 0, \\ & \mathcal{J}_k^i \leq \mathcal{J}_k^{i,\max}, \end{aligned} \tag{S11}$$

with the bounds $\mathcal{J}_k^{i,\max}$ encoding the medium and the reaction directions. Composition is recovered from the optimal weights as $\phi_l = w_l/\mu_C$.

### S6 Metabolic modes and the growth fraction

#### S6.1 Definition and extraction

A metabolic mode is one macrochemical equation: a charge- and element-balanced conversion summarising, per unit of biomass formed, the net substrates a species consumes and the products it excretes when its internal metabolism operates at steady state. Modes are obtained by single-organism flux balance analysis under a specified set of exchange bounds, after which the resulting exchange vector is divided by the biomass flux.

A species able to adopt several distinct strategies is represented by several modes, among which the community program is free to mix. The set of modes is an *inner* approximation of the species' true exchange space: any conical combination of extracted modes is achievable by the organism, so a coarser mode set can only under-represent what the community is able to do, never over-represent it.

#### S6.2 Modes that support little or no growth

The formulation requires every mode to be normalisable per unit of biomass formed, since this is what makes $s_{kl}$ constant and the balances linear in the weights (Section S4). Certain metabolic activities yield little or no growth when performed in isolation: examples are the over-fixation of nitrogen by organism $Q$ in the toy community, and pure product overflow in the syngas acetogen. Their unconstrained flux distributions lie close to a zero-growth state and cannot be normalised.

Such a mode is extracted by imposing a growth rate equal to a small fraction $f$ of the organism's maximum on the same medium, then dividing by that value. The resulting equation is a valid macrochemical conversion, but its coefficients are large, because the yield is expressed per unit of a small quantity of biomass. This is the origin of the large coefficients of mode $Q_2$ in the main text and of modes $A_4$ and $A_5$ in the syngas co-culture.

Non-growth-associated maintenance is set to zero during mode extraction. A constant ATP drain divided by the biomass flux introduces a term scaling as $1/\mu$, which would make the coefficients growth-rate dependent and the equation no longer a fixed conversion. Maintenance, where required, enters the community problem separately (Section S8).

#### S6.3 Sensitivity of the community optimum to $f$

The value of $f$ is not a fitted parameter and the community optimum does not depend on it. Table S2 reports the syngas community solved over a 25-fold range of $f$, at two hydrogen supplies and with and without an acetate feed. The maximum community growth rate, the predicted abundance of *C. kluyveri* and the identity and weight of every active mode are identical to five significant figures throughout.

Table S2: Insensitivity of the syngas community optimum to the growth fraction $f$ used to extract the near-growthless overflow modes $A_4$ and $A_5$. Supplies are in mmol gDW$^{-1}$h$^{-1}$ with carbon monoxide fixed at 10. Mode weights are given as fractions of $\sum_l w_l$. Values are identical across all five values of $f$; the overflow modes carry zero weight at every optimum, so the coefficients that depend on $f$ do not enter the solution.

| $f$ | **Acetate** | $H_2$ | $\mu_C$ **($h^{-1}$)** | **Active modes (weight fraction)** |
|---|---|---|---|---|
| 0.02–0.50 | 0 | 4 | 0.13703 | $A_1$ 0.56, $A_2$ 0.02, $A_3$ 0.36, $K_3$ 0.06 |
| | 0 | 12 | 0.19036 | $A_1$ 0.02, $A_2$ 0.42, $A_3$ 0.49, $K_3$ 0.08 |
| | 2 | 4 | 0.13730 | $A_1$ 0.57, $A_3$ 0.37, $K_3$ 0.06 |
| | 2 | 12 | 0.19881 | $A_1$ 0.10, $A_3$ 0.77, $K_3$ 0.13 |

Values tested: $f = 0.02, 0.05, 0.10, 0.20, 0.50$. All rows agree to the digits shown. Predicted *C. kluyveri* fractions are 0.0591, 0.0799, 0.0614 and 0.1272 respectively, likewise independent of $f$.

The reason is structural. The overflow modes are dominated: any flux distribution they represent can be achieved more efficiently by a conical combination of the growth-coupled modes, so the simplex optimum assigns them zero weight. They are retained in the library because they are required to span the exchange space under conditions where the growth-coupled modes are infeasible, but they do not participate in any optimum reported in the main text.

### S7 Model definitions

#### S7.1 Two-member cross-feeding community

Organisms $P$ and $Q$ are hypothetical, with small hand-built internal networks sharing ATP and NADH. $P$ alone can use glucose and $Q$ alone can fix $N_2$. The four macrochemical equations, one per mode, are given in Table S3; these are the equations drawn in Figure 1B of the main text.

Table S3: Macrochemical equations of the two-member community, per unit of biomass formed. $X_P$ and $X_Q$ denote one unit of biomass of $P$ and $Q$. ATP and NADH are balanced internally and do not appear. Mode $Q_2$ is extracted at a small imposed growth rate (Section S6.2), which is why its coefficients are large.

| **Mode** | **Strategy** | **Macrochemical equation** |
|---|---|---|
| $P_1$ | respire glucose | $1.43 \text{ glucose} + 0.20 \text{ NH}_3 \to X_P$ |
| $P_2$ | ferment to succinate | $2.05 \text{ glucose} + 0.20 \text{ NH}_3 \to X_P + 0.63 \text{ succinate}$ |
| $Q_1$ | self-sufficient | $1.86 \text{ succinate} + 0.10 \text{ N}_2 \to X_Q$ |
| $Q_2$ | over-fix and export N | $92.76 \text{ succinate} + 40.63 \text{ N}_2 \to X_Q + 81.05 \text{ NH}_3$ |

Only $P_2$ secretes succinate and only $Q_2$ secretes ammonia, so the mutualism is enforced by the balances rather than assumed. In the cross-feeding environment (glucose bounded at 1, $N_2$ at 5, ammonia and succinate not supplied) the program returns $\mu_C = 0.605\,\text{h}^{-1}$, reproducing the value $0.6047\,\text{h}^{-1}$ obtained for the same community by full-stoichiometry cFBA to within $5 \times 10^{-4}$.

#### S7.2 Anaerobic digestion community

Five species are each represented by a single macrochemical equation, involving nine external metabolites. The equations were taken from the accompanying stoichiometric theory, where they were derived from the known growth physiology of each organism. The signed coefficients $n_{kl}s_{kl}$ used are given in Table S4.

Table S4: Signed macrochemical coefficients of the anaerobic digestion community, per unit of biomass formed. Positive values denote production, negative values consumption. Blank entries are zero. Species abbreviations: cb, *Clostridium butyricum*; mm, *Methanococcus maripaludis*; dv, *Desulfovibrio vulgaris*; dm, *Desulfococcus multivorans*; mb, *Methanosarcina barkeri*.

| **Metabolite** | **cb** | **mm** | **dv** | **dm** | **mb** |
|---|---|---|---|---|---|
| glucose | −1.07 | | | | |
| ammonium | −0.20 | −0.20 | −0.20 | −0.20 | −0.20 |
| acetate | +0.409 | | −0.35 | | −12.945 |
| hydrogen | +2.937 | −34.1 | −9.0 | | |
| butyrate | +0.614 | | | −1.38 | |
| methane | | +7.754 | | | +12.45 |
| bicarbonate | +2.3 | −9.0 | −0.3 | +4.5 | +12.5 |
| bisulphide | | | +2.1 | +2.9 | |
| sulphate | | | −2.1 | −2.9 | |

Glucose, ammonium and sulphate are importable; acetate, hydrogen, butyrate, methane, bicarbonate and bisulphide are exportable. Acetate, hydrogen and butyrate are additionally cross-fed within the community.

#### S7.3 Syngas co-culture

The genome-scale reconstructions of *C. autoethanogenum* (iCLAU786) and *C. kluyveri* (iCKL708) comprise 2064 reactions and 1823 metabolites. Four corrections were applied to the deposited models before mode extraction (Section S7.4). The eight macrochemical modes obtained are listed in Table S5.

Table S5: Macrochemical modes of the syngas co-culture, per unit of biomass formed. $\mu$ is the growth rate at which the mode was extracted and $\mu_{\max}$ the organism's maximum on the same medium. Modes $A_4$ and $A_5$ are near-growthless overflow conversions extracted at $f = 0.05$ (Section S6.2); they carry zero weight at every optimum reported. Minor exchanges (phosphate, sulphate, $H_2S$, $NH_3$, $H^+$, $H_2O$) are omitted from the displayed equations for legibility but are present in the model.

| **Mode** | **Organism** | $\mu$ | $\mu/\mu_{\max}$ | **Principal conversion** |
|---|---|---|---|---|
| $A_1$ | acetogen | 0.1066 | 1.00 | $93.85\ CO \rightarrow X + 48.31\ CO_2 + 7.44$ acetate |
| $A_2$ | acetogen | 0.1657 | 1.00 | $60.33\ CO + 60.33\ H_2 \rightarrow X + 14.14$ acetate $+ 1.39\ CO_2$ |
| $A_3$ | acetogen | 0.1888 | 1.00 | $52.97\ CO + 78.05\ H_2 \rightarrow X + 11.15$ ethanol |
| $A_4$ | acetogen | 0.0094 | 0.05 | $1059\ CO + 1062\ H_2 \rightarrow X + 514.4$ acetate |
| $A_5$ | acetogen | 0.0094 | 0.05 | $1059\ CO + 2091\ H_2 \rightarrow X + 514.4$ ethanol |
| $K_1$ | elongator | 0.0535 | 1.00 | $186.8$ ethanol $+ 88.01\ CO_2 \rightarrow X + 70.03$ hexanoate |
| $K_2$ | elongator | 0.1137 | 1.00 | $87.93$ ethanol $+ 43.97$ acetate $+ 31.13\ CO_2 \rightarrow X + 42.24$ hexanoate |
| $K_3$ | elongator | 0.1475 | 1.00 | $67.82$ ethanol $+ 75.26$ acetate $+ 15.25\ CO_2 \rightarrow X + 43.32$ hexanoate |

Modes were defined by imposing, in turn, each substrate combination on which the organism can grow and each qualitatively distinct product it can form from that combination. For the acetogen these are carbon monoxide alone ($A_1$), carbon monoxide with limiting hydrogen ($A_2$), and carbon monoxide with hydrogen in excess, under which ethanol replaces acetate ($A_3$). For the chain elongator they are ethanol with carbon dioxide ($K_1$) and ethanol with acetate in balanced ($K_2$) or excess ($K_3$) proportion.

#### S7.4 Curation of the deposited reconstructions

Four defects in the deposited co-culture model were corrected. Each alters the growth or electron balance and would otherwise distort the extracted modes.

1. **Free cytosolic acetate uptake.** The reaction `EX_AC_c` admits acetate directly into the cytosol of *C. autoethanogenum*, bypassing the transport layer. Left open it inflates growth by approximately 65%. The reaction was closed.
2. **ATP maintenance bounds.** The deposited lower bounds (`ATPM_auto` 8.4, `ATPM` 0.45) are approximately tenfold the values used in the source publication (0.60 and 0.068) and render the model infeasible at the reported carbon monoxide uptake. They were set to the published values, and to zero during mode extraction (Section S6.2).
3. **Dissimilatory sulphate reduction.** As deposited, *C. kluyveri* reduces sulphate to $H_2S$ and excretes it, consuming three NADH per sulphate, so that sulphate acts as an electron sink in an organism that is a fermenter rather than a sulphate reducer. Blocking $H_2S$ excretion leaves

sulphate uptake free but restricts it to assimilation, after which uptake falls to the biomass sulphur demand.

4. **Hexanol production in *C. kluyveri*.** The hexanol route carries no gene–protein–reaction association and diverts electrons that otherwise yield hexanoate. Hexanol excretion was blocked in *C. kluyveri* only; butanol, which is gene-supported and carries no flux at the optimum, was left unmodified.

### S8 Extension: non-growth-associated maintenance

The results of the main text use the pure linear program (S11) and do not include non-growth-associated maintenance. This section records how the formulation accommodates it, for completeness.

Growth-associated maintenance adds ATP to the biomass reaction of each species and leaves the macrochemical coefficients growth-invariant, so (S11) is unchanged. Non-growth-associated maintenance is different: it is an ATP demand proportional to the biomass *present*, that is to $\phi_l$, whereas every other term scales with biomass present multiplied by growth, $w_l = \phi_l \mu_C$. Expressed in the weight variables it appears as a term proportional to $w_l/\mu_C$, which is non-convex and diverges as $\mu_C \to 0$.

The formulation remains tractable in absolute per-species biomass $X_l$, normalised so that $\sum_l X_l = 1$. Each external balance then carries a growth term and a maintenance term,

$$\sum_l (a_{lk}\,\mu_C + g_{lk}) X_l = b_k, \tag{S12}$$

where $a_{lk}$ is the macrochemical coefficient per unit biomass produced and $g_{lk}$ the maintenance exchange per unit biomass present. At fixed $\mu_C$ every coefficient is constant and (S12) is linear in the $X_l$. Feasibility is monotone in $\mu_C$, with a single crossover, so the maximum community growth rate in the presence of maintenance is obtained by bisection on that scalar, independently of the number of species. Per-biomass export caps, which are non-linear in the weight variables, are also linear in the $X_l$.

We note that this extension reintroduces the outer search over $\mu_C$ that (S10) removes. This is the price of a maintenance term, not a property of the reduction.

### S9 Software, data and reproducibility

Genome-scale models were handled with COBRApy. All linear programs were solved with a simplex method. The community programs for the two-member and anaerobic digestion communities were solved with the SciPy implementation of the simplex algorithm.

Code, models and the scripts that generate every figure and table in the main text and in this document are available at `https://github.com/TP-Watson/Community-FBA-from-metabolic-strategies/tree/main`. Exact package versions are recorded in the repository.

---

This document supports the main text; the macrochemical formulation, the worked examples and the ecological results are developed there.
